# *In silico* design of glyco-D,L-peptide antiviral molecules

**DOI:** 10.1101/583765

**Authors:** Luigi Agostini, Stefano Morosetti

## Abstract

**Background:** most licensed antiviral drugs are nucleoside analogs. A recent research focuses on blocking a virus from entering the cells in the viral cell adsorption/entry stage. In this entry mechanism the glycans present on the viral surface play a fundamental role. Homochiral L-peptides acting this fusion mechanism have shown some inhibition of viral infection. Peptides with regularly alternating enantiomeric sequence (L,D-peptides) can assume structures, which are not accessible to the corresponding homochiral molecules. Further, L,D-peptides are less sensitive to the enzymatic digestion.

**Aim:** *in silico* design a L,D-peptide with a high affinity for the viral surface glycans, and consequently able to interfere with its fusion mechanism.

**Methods:** a 3α,6α-Mannopentaose (3-6MP) molecule was used to simulate a viral surface glycan. Molecular Dynamics (MD) simulations of 3-6MP and D,L-peptide in water are performed using the force field AMBER12-GLYCAM06i. The binding constant was evaluated from trajectories. The D,L-peptide molecule was modified over the sequence, the length, the terminals and finally glycosylated to attain a very high binding constant value for 3-6MP.

In addition, the specific interaction between T lymphocyte CD4 glycoprotein and HIV envelope gp120 glycoprotein was studied through MD simulations between a D,L-peptide, bounded to a typical CD4 glycan, and a highly conserved HIV gp120 glycan.

**Results:** in the case of interaction with 3-6MP molecule, the very effective molecule obtained was H-D-Trp-L-Pro-D-Asn-L-Pro-D-Trp-L-Pro-D-Asn-L-Pro-OH where the Asn residues in position 3 and 7 are glycosylated with alpha-D-Mannopyranosyl-(1->3)-[alpha-D-mannopyranosyl-(1->6)]-alpha-D-mannopyranosyl-(1->4)-N-acetyl-beta-D-glucopyranosyl-(1->4)-N-acetyl-beta-D-glucopyranosyl-1-OH.

In the case of interaction with HIV envelope, the very effective molecule obtained, able to antagonize the CD4 glycoprotein, was H-D-Trp-L-Pro-D-Asn-L-Pro-D-Trp-L-Pro-D-Asn-L-Pro-OH where the Asn residue in position 3 is glycosylated with alpha-D-galactopyranosyl-(1->4)-N-acetyl-beta-D-glucopyranosyl-(1->2)-alpha-D-Mannopyranosyl-(1->3)-[alpha-D-galactopyranosyl-(1->4)-N-acetyl-beta-D-glucopyranosyl-(1->2)-alpha-D-Mannopyranosyl-(1->6)]-beta-D-mannopyranosyl-(1->4)-N-acetyl-beta-D-glucopyranosyl-(1->4)-N-acetyl-beta-D-glucopyranosyl-1-OH

**Conclusion:** these glycosylated D,L-peptide molecules are very promising representatives of a new class of antiviral agents.

## Introduction

There is a continuous effort in the search for new antiviral agents, which aims to develop new and more effective drugs, capable of targeting an increasing number of diseases while, at the same time, limiting their side effects. The approved drugs have different mechanism of actions during the viral life cycle^1^. A few of them are entry inhibitors. One of these (Enfuvirtide) is a polypeptide with 36 units. Many polypeptides were tested as antiviral^2,3^. The viral fusion stage is often mediated by the glycoproteins present on its envelope^4^.

The oligo-L,D-peptides present structures which are not accessible to the corresponding homochiral peptides. In particular, by assuming local β conformations, they can realize helices of different diameter^5,6^. Another property of these molecules with the peptide bond connecting D,L and L,D dimers is the decreased susceptibility to digestion by proteolytic enzymes compared with the LL dimers^7^. This results in their increased average life time in the organism.

The initial idea of our research was to find a short L,D-oligopeptide capable of binding the sugar moiety of the viral surface glycoproteins, so interfering with the fusion stage of the infection. For this purpose we performed different *in silico* Molecular Dynamics simulations (MD) on a system consisting of a L,D-peptide molecule and a oligosaccharide.

The chosen oligosaccharide is 3α,6α-Mannopentaose (3-6MP) (alpha-D-Mannopyranosyl-(1->3)-[alpha-D-mannopyranosyl-(1->3)-[alpha-D-mannopyranosyl-(1->6)]-alpha-D-mannopyranosyl-(1->6)]-beta-D-mannopyranose) (fig.1). This is a common core structure for a large number of glycans present in glycoproteins^8^.

**Fig. 1.**
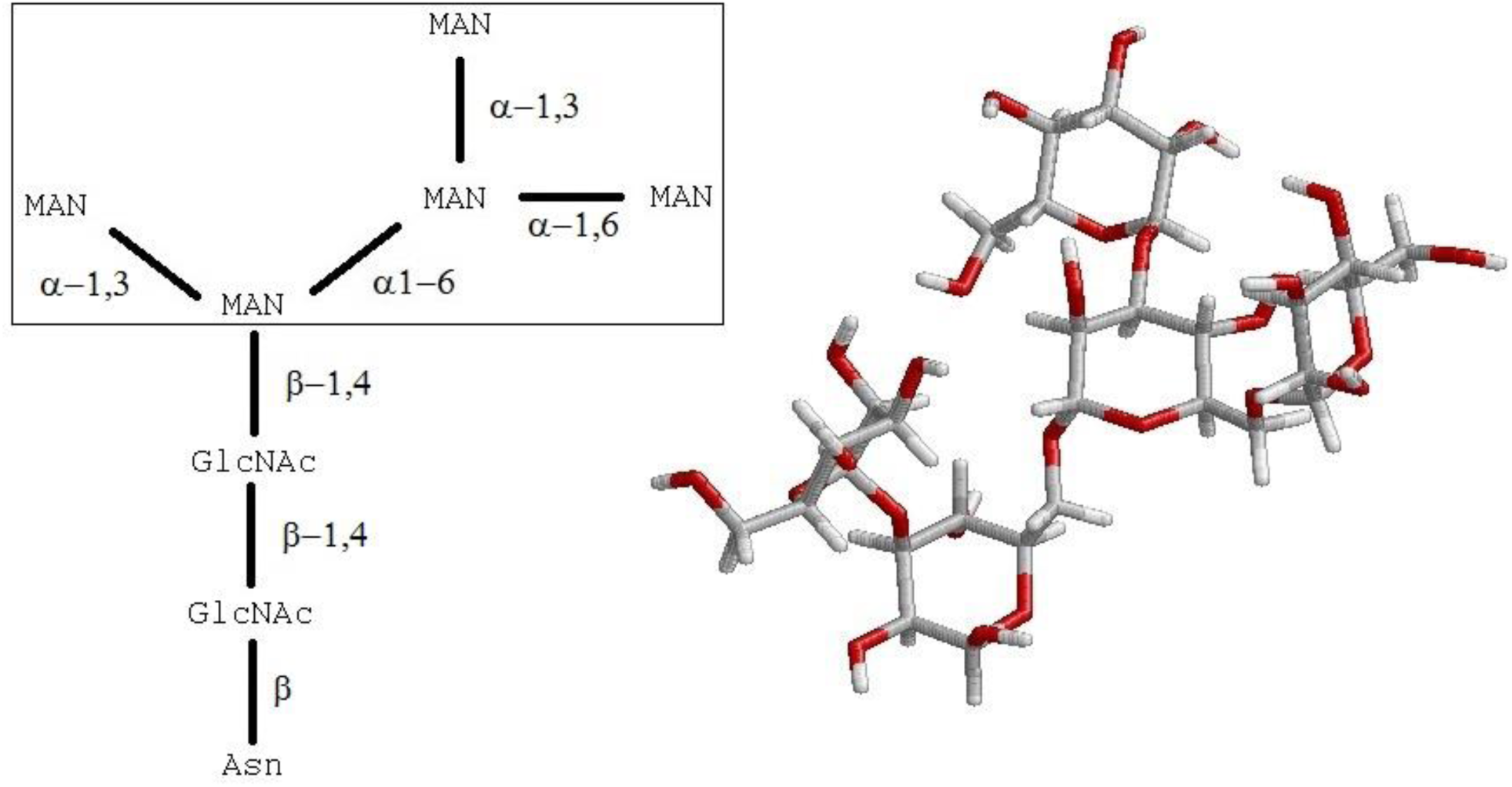
Right: the 3-6MP molecule representing the glycan component of the glycoproteins. Left: The schematic representation of the molecule is enclosed in a rectangle. Outside the square the link with the protein portion is shown.

The D,L-peptide molecule initially chosen for the simulation is H-(D-Phe-L-Lys)_4_-OH. The octapeptide can assume a β-helix conformation with 6.2 monomer per turn^6^. Our initial guess was that the helix would turn around the oligosaccharide (fig.2).

**Fig. 2.**
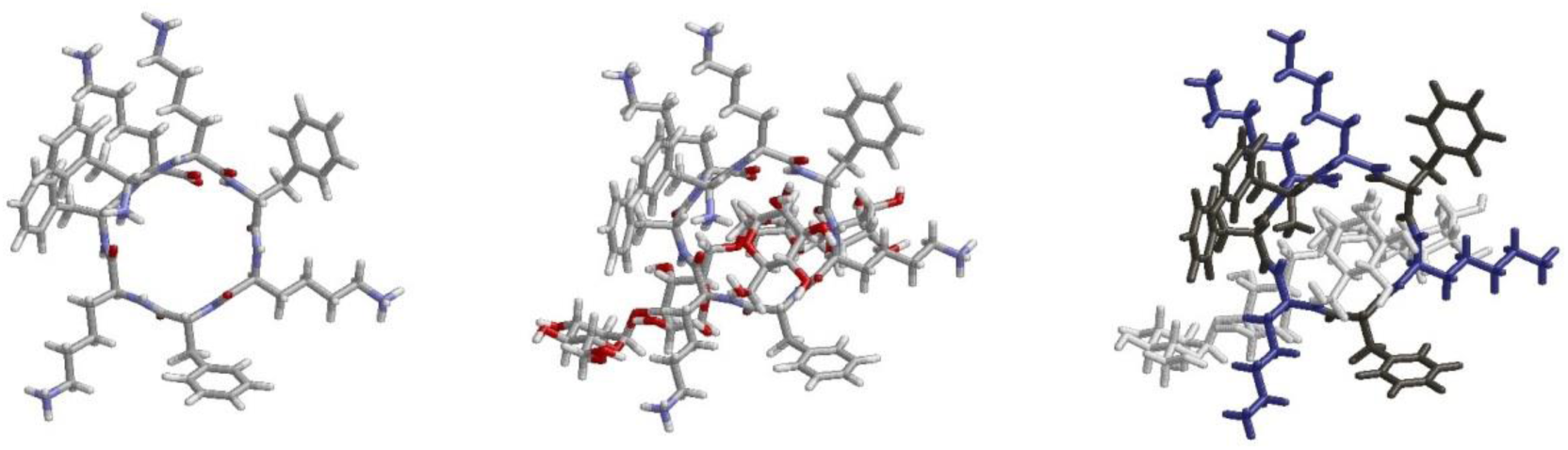
Left: H-(d-Phe-L-Lys)_4_-OH in β-helix conformation with 6.2 monomer per turn. Middle: the helix turns round the oligosaccharide. Right: same as the middle, but with different colors for the oligosaccharide, Phe residues, Lys residues.

The rationale for the octapeptide chosen sequence was its solubility, obtained through the introduction of the charged Lys residues, and the stabilizing van der Waals interactions between Phe residues and the oligosaccharide.

The association of the two molecules was monitored performing MD simulations, through visual inspection of the trajectory by means of VMD 1.9.1^9^ program and following the molecules’ center of gravity distance as a function of time.

Our simulations, however, showed that this system (fig.2) is unstable. The peptide molecule moves away from the oligosaccharide and evolves into a globular structure, where β-turns are present (fig. 3). This happens because the peptide sequences L,D and D,L can easily form β-turn conformations^10^. Therefore, the resulting association constant has a low value.

**Fig. 3.**
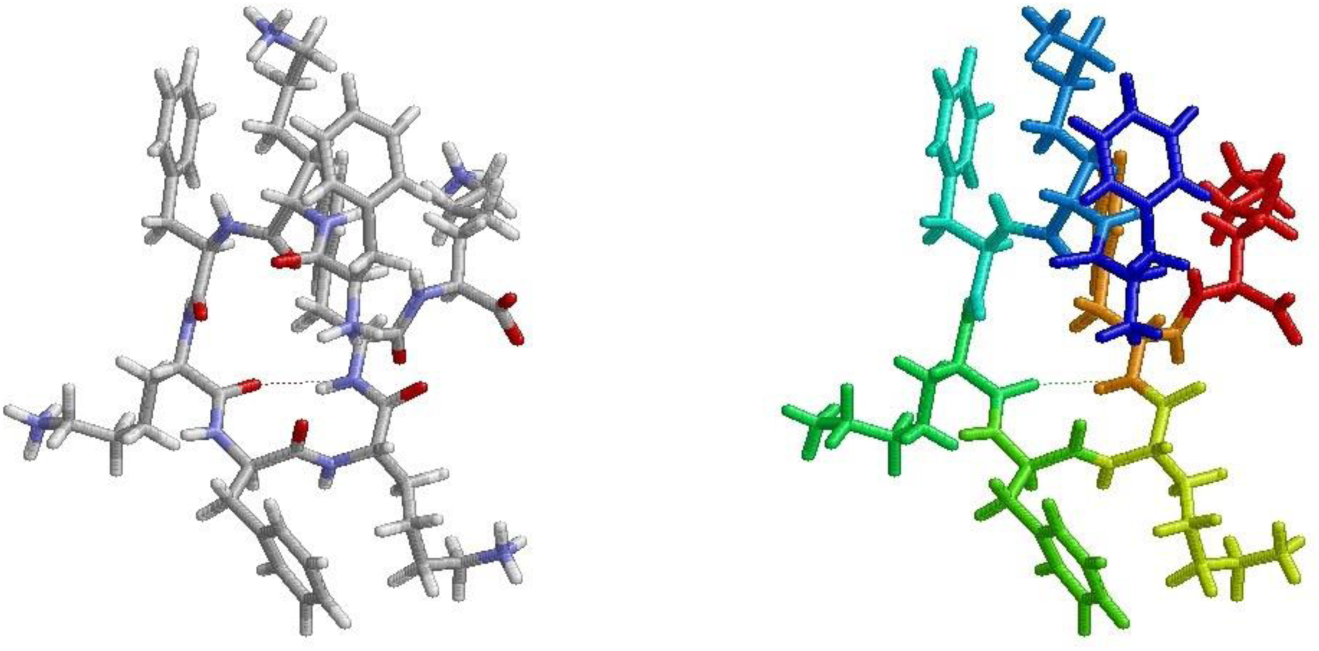
Globular conformation, with hydrogen bond highlighted by dashed line in the β-turn. Left: CPK colors, right: peptide groups highlighted by different colors. The conformation is taken from a MD trajectory.

Given this result, we decided to modify the D,L-peptide molecule over the sequence, the length, the terminals and finally glycosylated in an attempt to attain a very high binding constant value. It is worth mentioning that glycopeptide antibiotic molecules have shown antiviral activity^11,12^. Detailed Methods and Results are given below.

## Methods

### Force field

oligosaccharides are present in the system with the peptide molecule, hence the choice for the force field is AMBER12sb^13^ with GLYCAM06j^14,15^.

### Molecular Dynamics simulations

All-atom MD simulations were performed in water by using GROMACS 4.5.6 software package^16^. AMBER12sb and GLYCAM06j was obtained from the pertinent official sites^17,18^. After minimization and 150 ps equilibration steps on a *NVT* ensemble and subsequent 150 ps equilibration steps on *NPT* ensemble, a 200 ns MD simulation was performed in a periodic box, with 1 Na^+^ and 1 Cl^-^ to simulate the ionic strength.

The following settings were used: An integration step of 2 fs with periodic boundary conditions at constant volume and temperature (T=310 K). Leap-frog integration scheme of the equations of motion. The default LINCS algorithm to constrain bonds involving H atoms. The Verlet cutoff scheme for neighbor searching, since parallel computing on GPUs was performed. Particle Mesh Ewald (PME) algorithm for long range electrostatics.

Three different simulations were performed for each peptide molecule, differing in box volume and/or relative position of peptide and 3-6MP molecules and/or initial peptide conformation.

### Carbohydrate coordinates, topology and charges

they were obtained by the Carbohydrated Builder tool present in the GLYCAM-web site glycam.org. The molecule can be submitted in condensed Glycam notation^19^ or can be built specifying the monosaccharides and their linkages. Files with the extensions rst7 for coordinates and parm7 for topology were obtained, in AMBER style.

The program glycam2gmx.pl^20,21^ can be used to obtain files in GROMOS stile. The command line is perl ./glycam2gmx.pl –prmtop *.parm7 –crd *.rst7 –outname *, which returns *.gro for coordinates and *.top for topology (* stands for the chosen names).

These files must be merged with the corresponding peptide.gro and peptide.top files obtained by the program pdb2gmx for the peptide molecule.

One coordinates file was obtained merging the carbohydrate coordinates in the peptide.gro file. They can be visualized by VMD program and translated through the program editconf (present in the GROMACS suite) if necessary.

One topology file was obtained by renaming the carbohydrate file *.top to *.itp and inserting the line #include “*.itp” in the peptide.top file. Duplicated records, atom types definitions and water topology must be deleted.

### Glycopeptide coordinates, topology and charges

they were obtained by the Glycoprotein Builder tool present in the GLYCAM-web site glycam.org. Step 1: the coordinates of the peptide under glycosylation are submitted in pdb^22^ format, without hydrogens and records CONNECT. Step 2: the ends are fixed. If the peptide is in zwitterionic form, CONTINUE must be chosen. Step 3: the glycan derivative is chosen by the interactive carbohydrated builder tool; the saccharide chain is selected, ending with aglycon –OH; this end will be linked to the peptide; after aglycon, branches can be introduced; term with DONE command; add glycan to glycosylation sites: N-linking and O-linking biologically likely sites are shown; SHOW ALL gives all sites with the solvent accessible surface area; select and continue; download current structure. Step 4: the structure is minimized. Step 5: download of the rst7 and parm7 files.

The program glycam2gmx.pl can be used to obtain files in GROMOS stile. In simulations of the glycopeptide with a carbohydrate molecule, its coordinates and topology will be merged in glycopeptide files by following the same procedure as before.

### Association constant estimation

the distance between the molecules’ centers of gravity was calculated by the command g_dist for each MD simulation. When the intermolecular distance was lower than a certain value dependent upon the molecular weights (MW) of the molecules (in our case 1.5 nm) these molecules were considered associated. From the total time of the MD simulation (t_tot_) is possible to extract a time when the molecules are considered associated (t_ass_). Through the introduction of *f* = *t*_*ass*_/*t*_*tot*_, the fraction of association time, the association constant *K*_*ass*_ can be roughly calculated as:

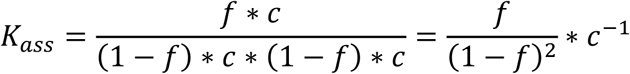

where the molar fraction is considered numerically equal to the time fraction, applying the ergodic postulate. The trajectories are of limited extension, and hence the approximation in constant evaluation.

The association constants reported were averaged over all trajectories calculated for the same molecular system. The error was calculated as half the range of values obtained.

## Results

The peptide or glycopeptide sequences studied are given below together with their *K*_*ass*_ (in mol^-1^) with 3-6MP molecule.

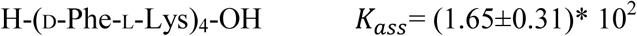

The distance vs time between the peptide and 3-6MP molecules in a single trajectory, is shown in fig. 4.

**Fig. 4.**
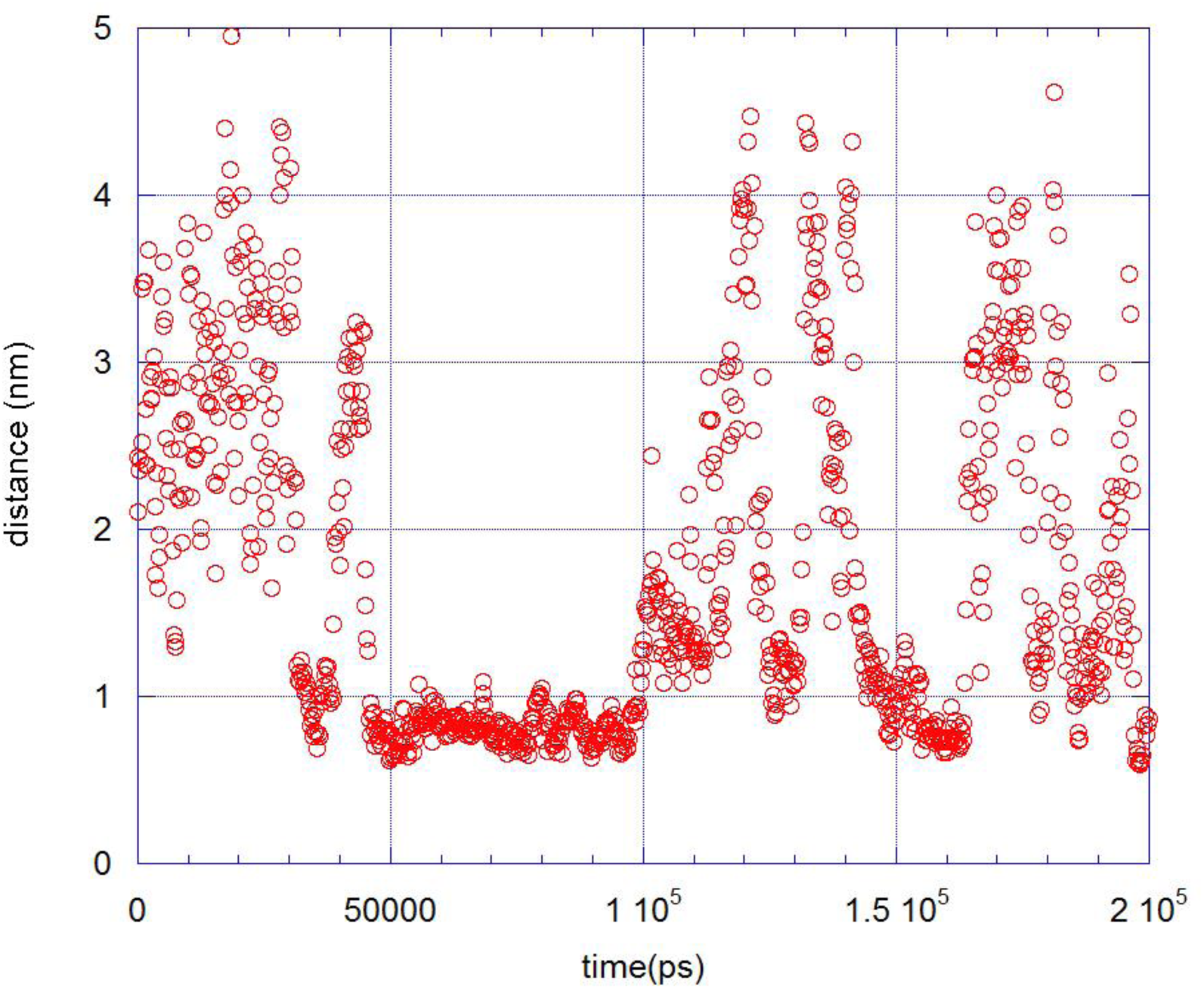
Distance (nm) between H-(d-Phe-L-Lys)_4_-OH and 3-6MP molecules, reported every 200 ps; the total time is 200 ns.

The effect of changing the terminal and amino acid sequence was explored with the following MD simulations, where FOR stands for the formilated NH_2_, and Hyp is the hydroxyproline amino acid:

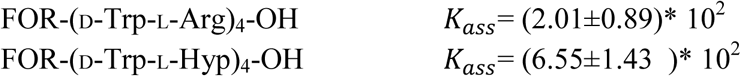

The presence of the Hyp seems to have a moderate effect on increasing the binding costant.

In order to achieve the expected association constant, we introduced the following changes: 1) terminal ACE and NME were introduced to increase the number of intramolecular hydrogen bonds (HB) so as to stabilize the peptide helix conformation; 2) Pro or Hyp were introduced to impart conformational stiffness to the helix; 3) The D-aminoacid was replaced to try to introduce a range of hydrophobic and hydrogen bonding interactions.

All the following peptides behave like the peptide H-(D-Phe-L-Lys)_4_-OH in spite of the changes introduced, as evidenced by the trajectory displayed and the *K*_*ass*_ obtained.

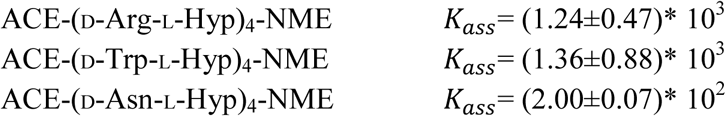

The effect of the peptide’s length was also studied through the following sequences:

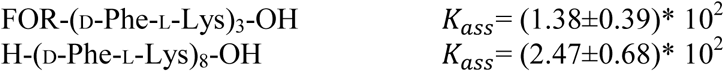

Finally, a peptide with an overall negative charge, instead of positive, was studied:

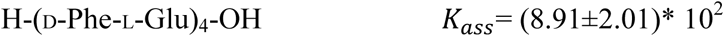

All sequences shown do not exhibit a *K*_*ass*_ much higher than the initial peptide. The introduction of a amino acid D-Trp, D-Arg or the rigid conformation amino acid L-Hyp, increased the *K*_*ass*_ by a little less than an order of magnitude.

Peptide glycosylation was introduced. The further interactions with 3-6MP molecule, were expected to increase the *K*_*ass*_.

H-D-Asn-L-Pro-D-Asn(-GlcNAc)-L-Pro-D-Asn(-GlcNAc)-L-Pro-D-Asn(-GlcNAc)-L-Pro-OH

*K*_*ass*_= (1.11±0.67)* 10^3^

GlcNAc stands for N-acetyl-glucosamine.

In consideration of these disappointing results, a more complex glycosylation was tried. The glycoside attached to the peptide was: alpha-D-Mannopyranosyl-(1->3)-[alpha-D-mannopyranosyl-(1->6)]-beta-D-mannopyranosyl-(1->4)-N-acetyl-beta-D-glucopyranosyl-(1->4)-N-acetyl-beta-D-glucopyranosyl-1-OH (abbreviated as Penta). See fig. 5. It is the common motif in the high mannose and complex N-linked glycan chains of the glycoproteins.

**Fig. 5.**
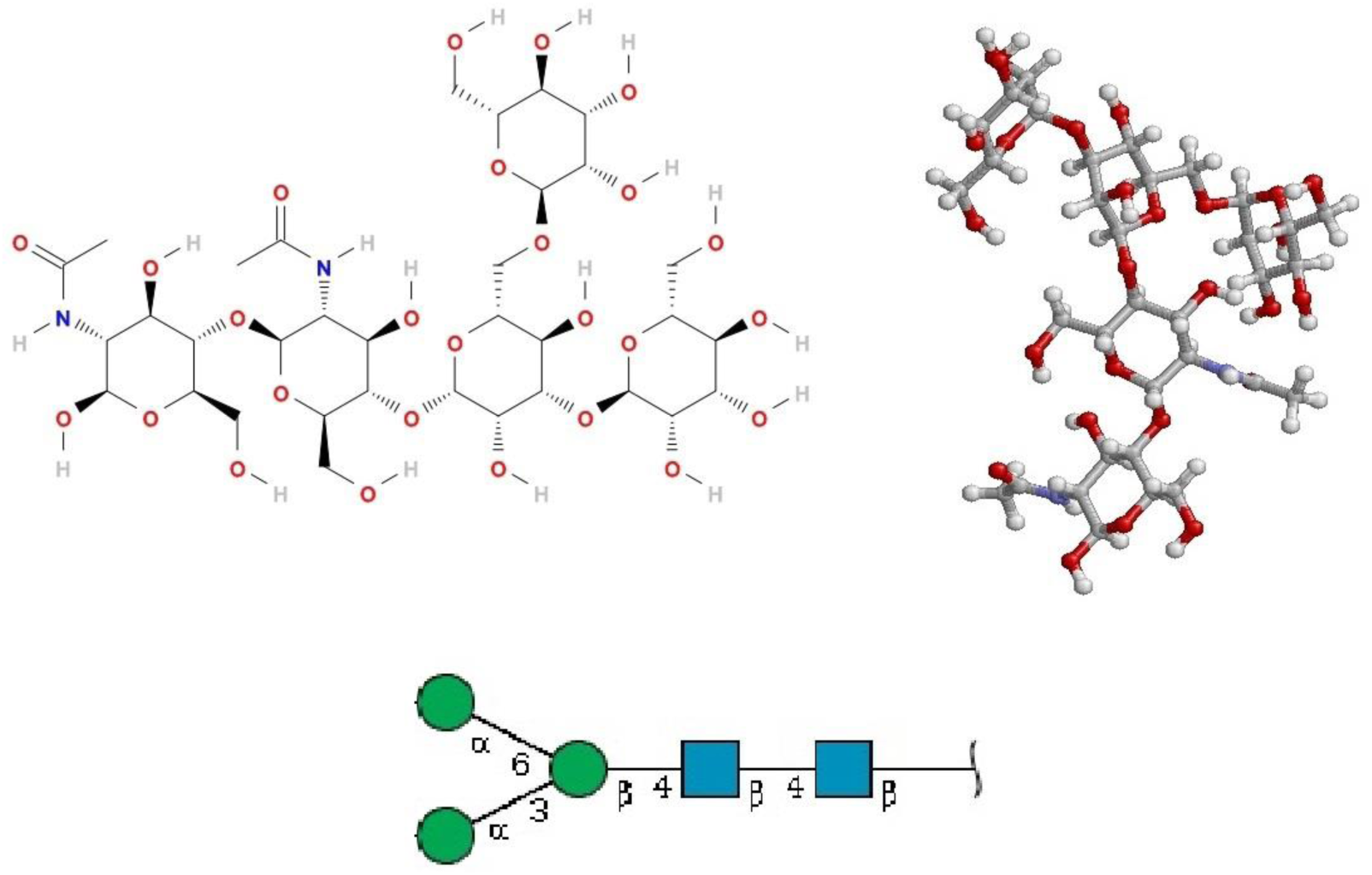
The glycoside Penta attached to peptide. Upper left: scheme. Upper right: structure. Bottom: SNFG representation^23^.

The molecule submitted to MD simulation was H-D-Trp-L-Pro-D-Asn(-Penta)-L-Pro-D-Trp-L-Pro-D-Asn(-Penta)-L-Pro-OH (abbreviated as TrpProAsnPenta). Where: Pro is present for its rigidity and Trp was introduced because of the higher *K*_*ass*_ obtained when present in the previously studied molecules. The molecule is shown in fig. 6,

**Fig. 6.**
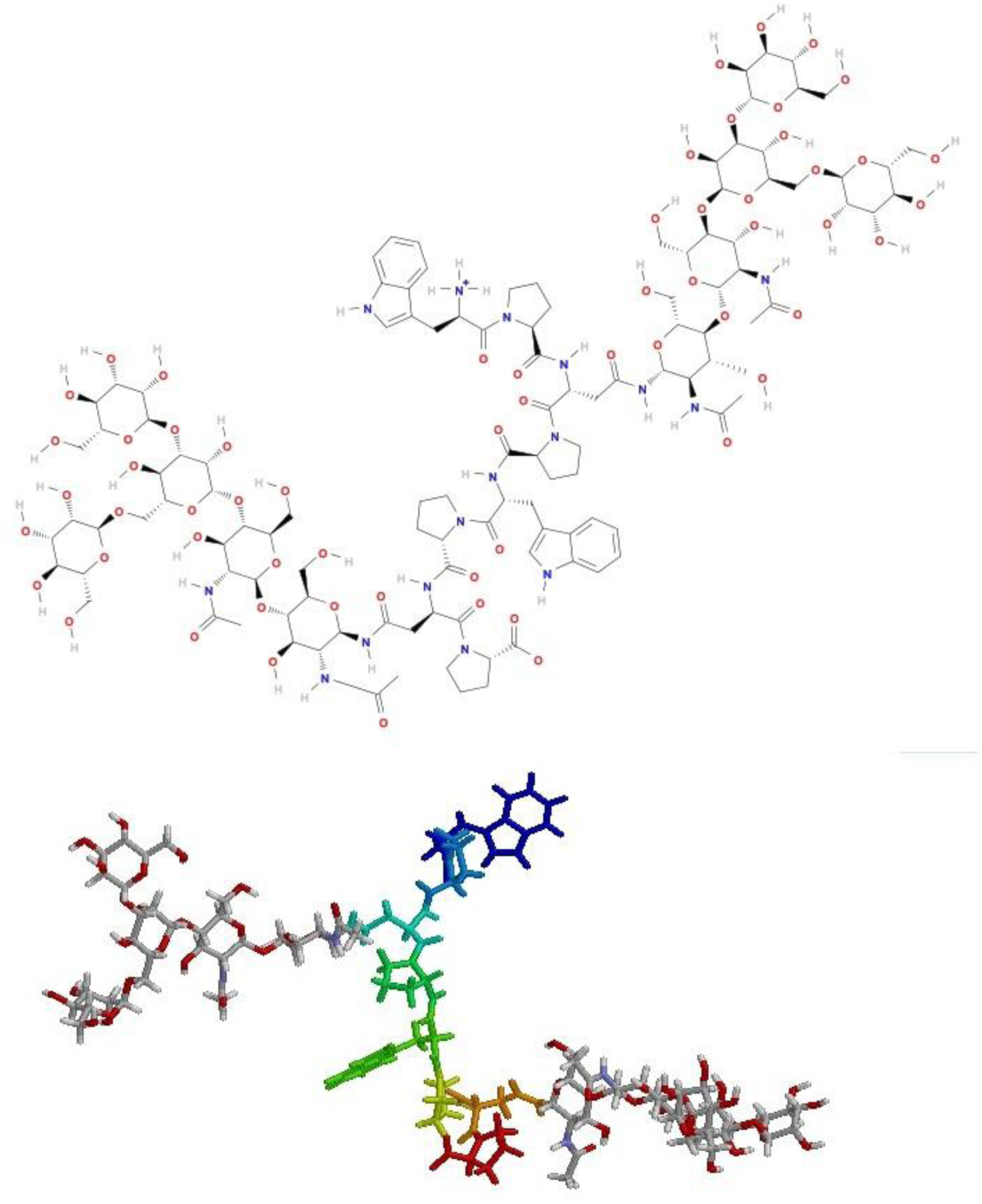
H-D-Trp-L-Pro-D-Asn(-Penta)-L-Pro-D-Trp-L-Pro-D-Asn(-Penta)-L-Pro-OH. Up: scheme. Down: structure; the oligopeptide is shown in elongated structure and the single aminoacids in different colors for clarity.

In this case six different MD simulations were performed. The results are in all cases a stable bending between TrpProAsnPenta and 3-6MP after their initial matching into the simulation box. There can be different arrangements in the bending, but without dissociation. This demonstrates a very high *K*_*ass*_ (not measurable with the aforementioned method).

The results of a MD simulation are reported as the inter-molecular distance vs time (fig.7) and in (fig.8) is reported the structure of the association complex obtained.

**Fig. 7.**
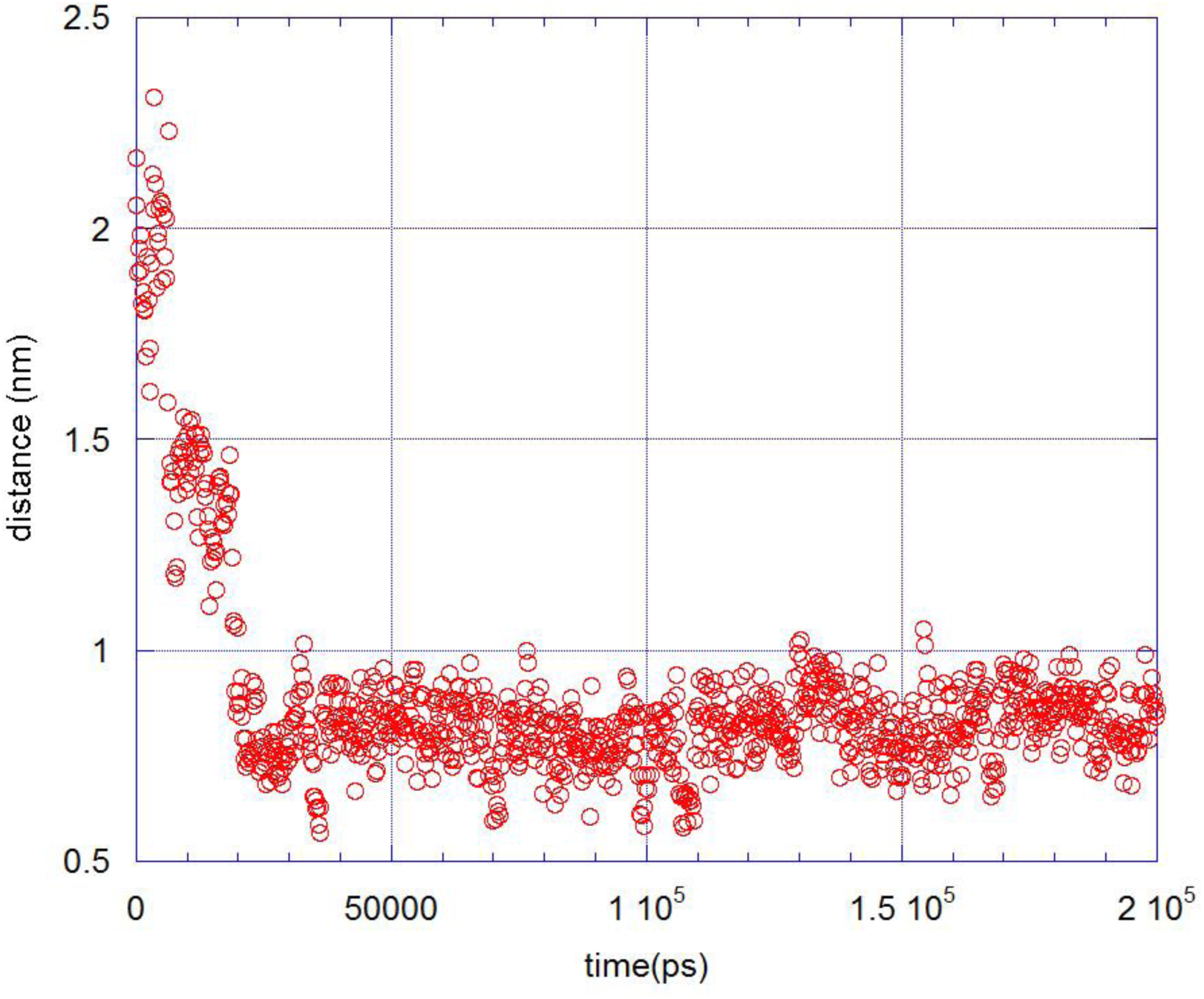
Distance (nm) between TrpProAsnPenta and 3-6MP molecules, reported every 200 ps; the total time is 200 ns.

Encouraged by these positive findings, we developed a simulation aimed to a specific case. The idea is to use for glycosylation the same cellular glycan chain preferentially exploited by the virus for the adhesion in the first stage of fusion, thus creating a molecular mimicry against the virus. Considering the specific case of HIV virus, this means using a glycan chain of the CD4 glycoprotein, which is involved in the interaction with the viral surface glycoproteins in the early stage of recognition^4^. Numerous different glycan chains are reported for this glycoprotein^24^, with respectively: one, two or three branches. We chose the structure alpha-D-galactopyranosyl-(1->4)-N-acetyl-beta-D-glucopyranosyl-(1->2)-alpha-D-Mannopyranosyl-(1->3)-[alpha-D-galactopyranosyl-(1->4)-N-acetyl-beta-D-glucopyranosyl-(1->2)-alpha-D-Mannopyranosyl-(1->6)]-beta-D-mannopyranosyl-(1->4)-N-acetyl-beta-D-glucopyranosyl-(1->4)-N-acetyl-beta-D-glucopyranosyl-1-OH (abbreviated as 9glyco) reported in fig. 9 as our first representative, because it has the most recurrent monosaccharides for each given position, in the most common two branched motif.

**Fig. 8.**
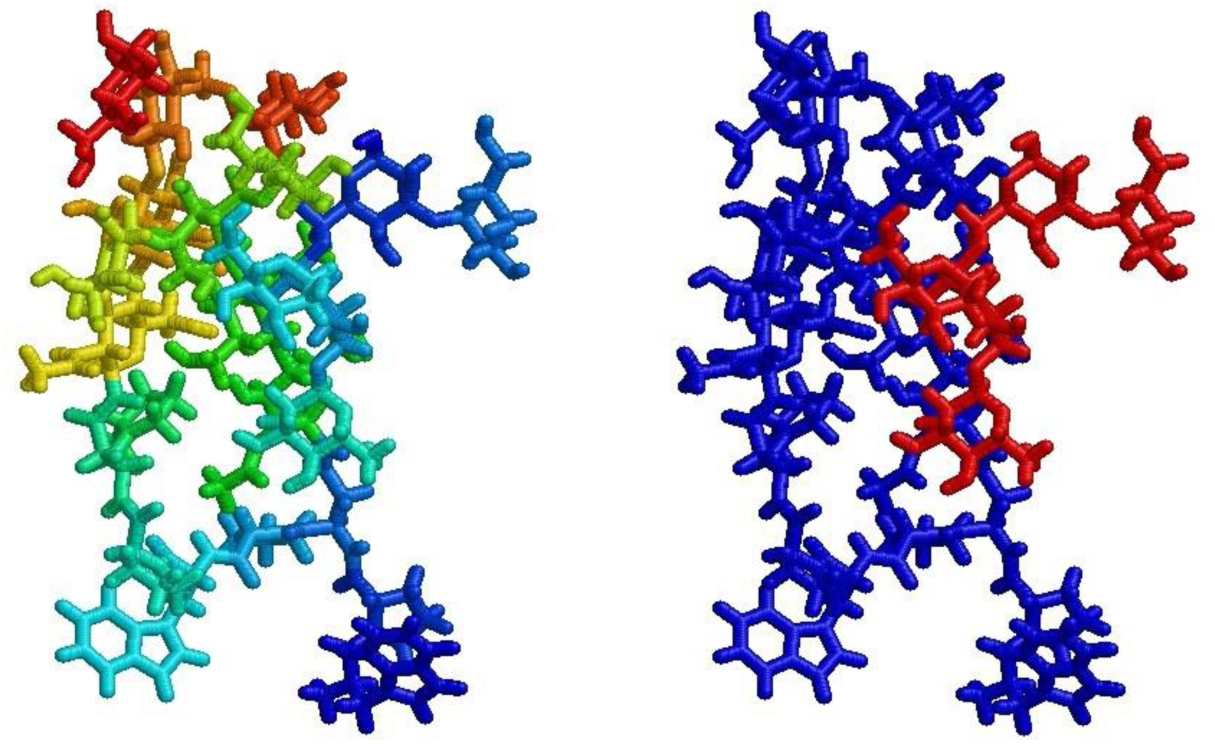
85 ns time structure, which represents the average of the association during the simulation; two representations are shown, one with different colors for different chemical units, the other with different colors for the 2 molecules, to locate them better.

**Fig. 9.**
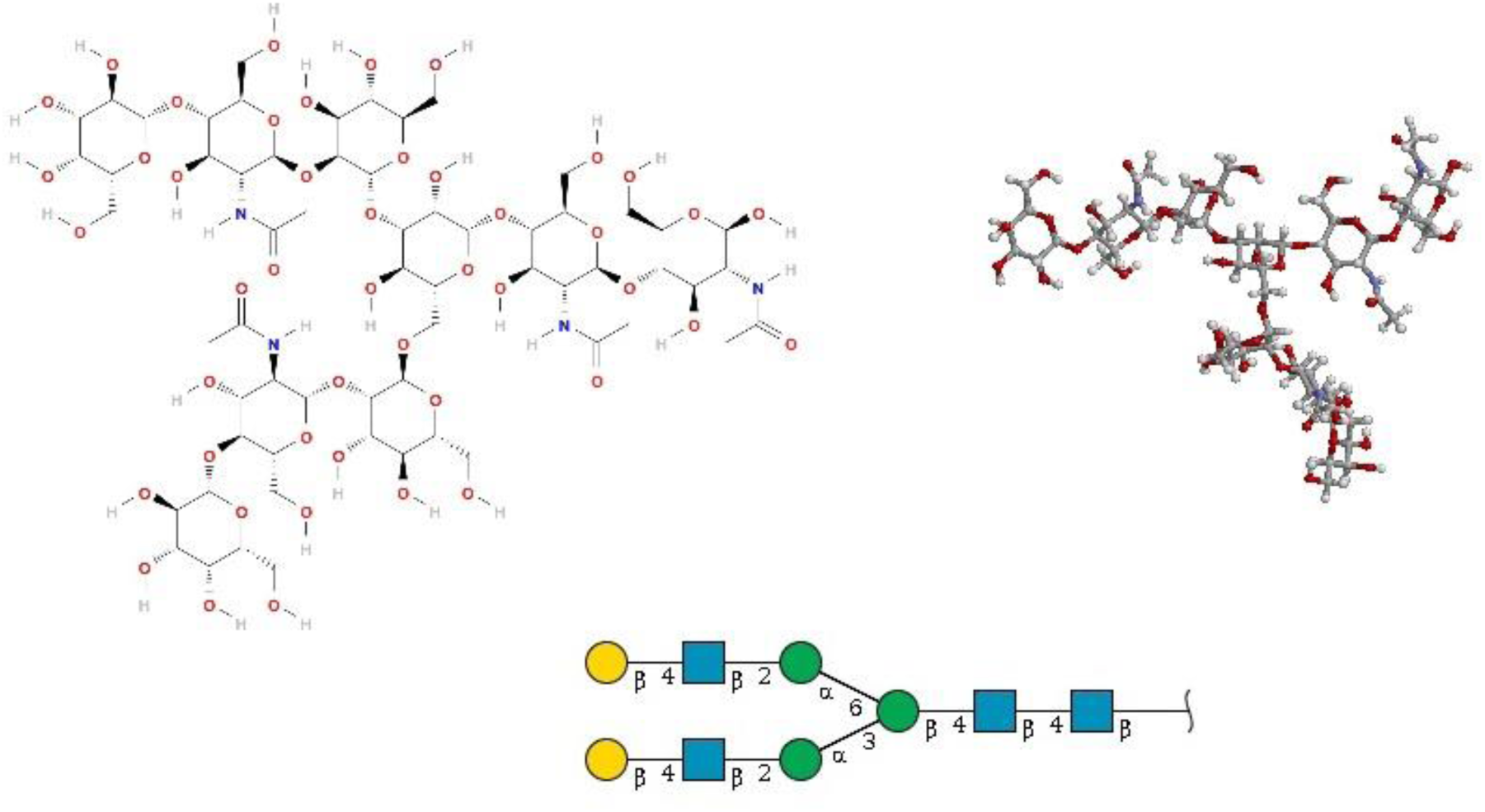
The glycoside 9glyco attached to peptide. Upper left: scheme. Upper right: structure. Bottom: SNFG representation^23^.

Consequently, the glycopeptide used in the simulations was H-D-Trp-L-Pro-D-Asn(-9glyco)-L-Pro-D-Trp-L-Pro-D-Asn-L-Pro-OH (abbreviated as TrpProAsn9glyco).

The glycoprotein CD4 interacts with the HIV glycoprotein gp120, which can relate through the exposure of different glycans^4^. The glycan at position Asn262 in HIV gp120 is chosen for the interaction with TrpProAsn9glyco, because it has the best resolved structure in a recent quaternary complex with CD4, and its presence is essential for the fusion mechanism^25^. The high mannose glycan used in the simulations is alpha-D-Mannopyranosyl-(1->2)-alpha-D-Mannopyranosyl-(1->2)-alpha-D-Mannopyranosyl-(1->3)-[alpha-D-Mannopyranosyl-(1->6)-[alpha-D-Mannopyranosyl-(1->3)**]-**alpha-D-Mannopyranosyl-(1->6)]-beta-D-mannopyranosyl-(1->4)-N-acetyl-beta-D-glucopyranosyl-(1->4)**-**N-acetyl-beta-D-glucopyranosyl-1-OH with the OH replacing the connection to Asn262 position. It is here abbreviated as N262glyco and reported in fig.10.

**Fig. 10.**
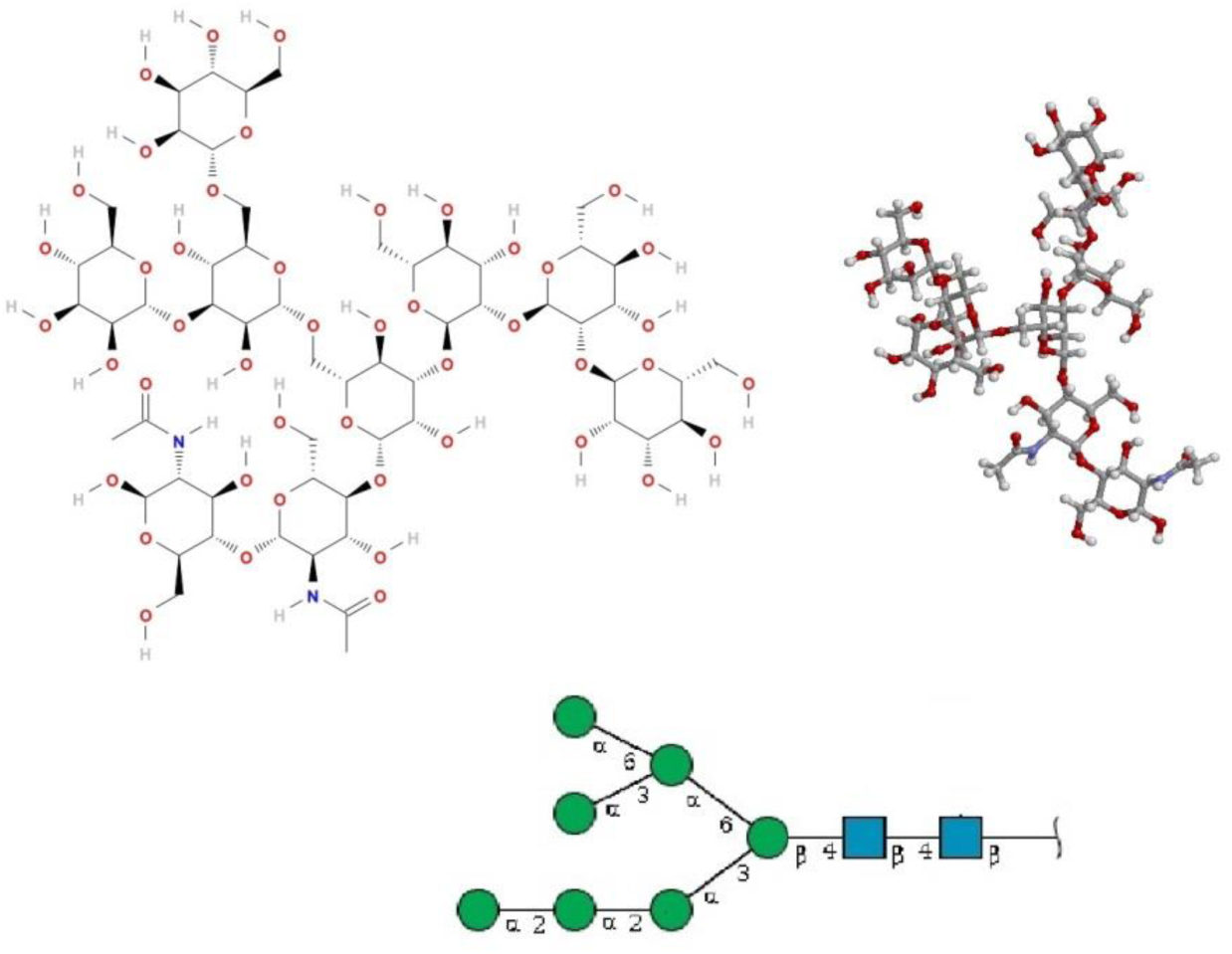
The glycoside N262glyco. Upper left: scheme. Upper right: structure. Bottom: SNFG representation^23^.

MD simulations were performed over TrpProAsn9glyco and N262glyco in water. The results are in all cases a stable bending between TrpProAsn9glyco and N262glyco after their initial matching into the simulation box. Similar to the previous case, here too, we found a very high *K*_*ass*_ (not measurable with the aforementioned method).

The results of one MD simulation are reported (fig.11) showing the structures of the association complex obtained:

**Fig. 11.**
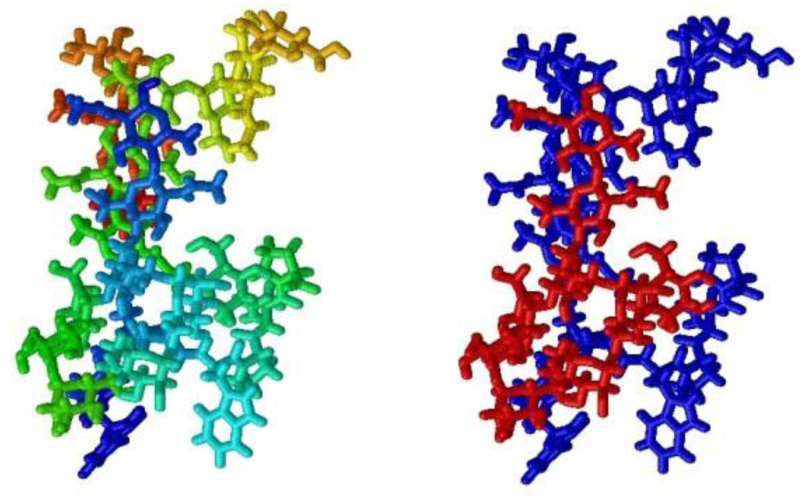
73.8 ns time structure, which represents the average of the association during the simulation; two representations are shown, one with different colors for different chemical units, the other with different colors for the 2 molecules, to locate them better.

It is worth mentioning that the molecules’ relative positions, in the ‘association state’, are consistent with a hypothetical interaction between CD4 and gp120 glycoproteins, as indicated by the opposite positions of the glycosides binding points, which are the peptide and the GlcNAc terminals, respectively. These findings are illustrated in fig. 12.

**Fig. 12.**
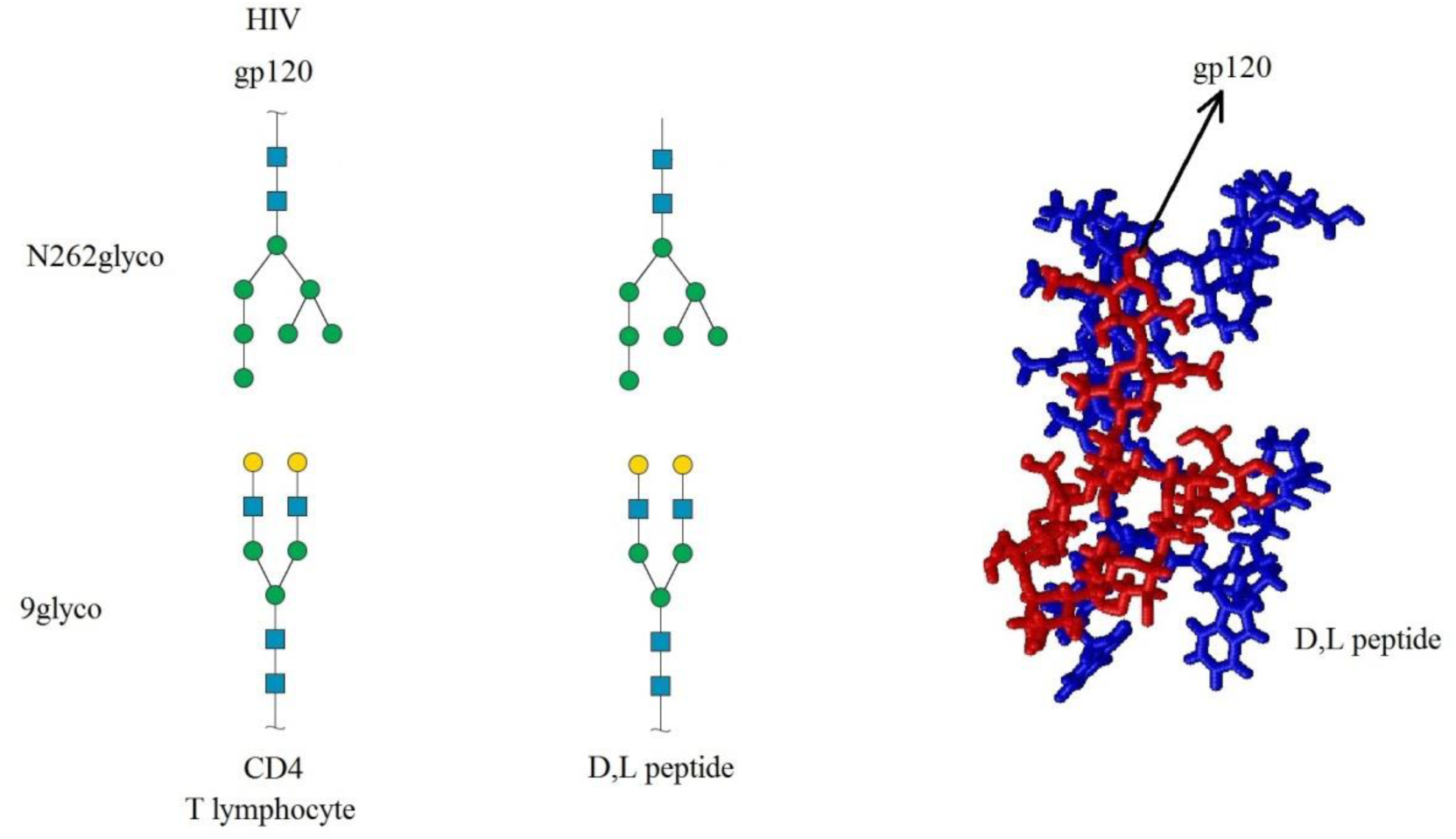
Interaction between the glycans 9glyco and N262glyco. Left: in vivo; the glycoproteins involved are indicated. Center: model interaction; the binding of 9glyco to D,L peptide is indicated. Right: the representative structure of the association obtained through MD simulation; red: N262glyco, blu: D,L peptide 9glyco. The position of D,L peptide is highlighted. The arrow indicates the GlcNAc bonded to gp120 in vivo.

## Conclusions

Our results showed that oligo-D,L-peptides can achieve a stable association with a glycan, only when glycosylated. Further, the glycosylation is effective only if the glycan side chains are suitably long.

Actually H-D-Asn-L-Pro-D-Asn(-GlcNAc)-L-Pro-D-Asn(-GlcNAc)-L-Pro-D-Asn(-GlcNAc)-L-Pro-OH which has three monomeric glycosylation sites, does not achieve a stable association with 3-6MP molecule.

H-D-Trp-L-Pro-D-Asn(-Penta)-L-Pro-D-Trp-L-Pro-D-Asn(-Penta)-L-Pro-OH realizes *in silico* the goal of a high bending with the molecule 3-6MP, which is the common core structure of a large number of glycans present in glycoproteins.

H-D-Trp-L-Pro-D-Asn(-9glyco)-L-Pro-D-Trp-L-Pro-D-Asn-L-Pro-OH realizes *in silico* the goal of a high bending with the molecule N262glyco, where 9glyco is a common glycosylation of CD4 and N262glyco a conserved glycan chain of gp120, so to realize a molecular mimicry against the HIV virus.

If our *in silico* studies will be confirmed by *in vivo* tests, this array of compounds will represent an entire new class of anti-viral drugs, with promising therapeutic potential.

A possible synthetic route for these molecules is presented in Appendix.

The great variability, achievable through modifications of the peptide sequence, length, endings and through its glycosylation, allows the possibility of fine tuning logP, solubility and interaction specificity.

Finally, it is possible to use for glycosylation the same cellular glycan chain preferentially exploited by the virus for the adhesion in the first stage of fusion, thus creating a molecular mimicry against the virus. In the case of HIV virus, we have used a glycosylic chain of CD4 glycoprotein^4^, involved in interaction with the surface glycoproteins of the virus in the early stage of recognition.

In future work, it will be extremely interesting to test less common glycosylations such as Arginine N-glycosylation^26^ and Arginine N-rhamnosylation^27^.

## Supporting information

figure 8 in rotation

figure 11 in rotation

## Abbreviations

3-6 MP: alpha-D-Mannopyranosyl-(1->3)-[alpha-D-mannopyranosyl-(1->3)-[alpha-D-mannopyranosyl-(1->6)]-alpha-D-mannopyranosyl-(1->6)]-beta-D-mannopyranose 9glyco: alpha-D-galactopyranosyl-(1->4)-N-acetyl-beta-D-glucopyranosyl-(1->2)-alpha-D-Mannopyranosyl-(1->3)-[alpha-D-galactopyranosyl-(1->4)-N-acetyl-beta-D-glucopyranosyl-(1->2)-alpha-D-Mannopyranosyl-(1->6)]-beta-D-mannopyranosyl-(1->4)-N-acetyl-beta-D-glucopyranosyl-(1->4)-N-acetyl-beta-D-glucopyranosyl-1-OH
CPK: atomic coloring scheme
FOR: formilated
GlcNAc: N-acetyl-glucosamine
GPU: Graphics Processing Unit
HB: hydrogen bond
MD: molecular dynamics
MW: molecular weight
N262glyco: alpha-D-Mannopyranosyl-(1->2)-alpha-D-Mannopyranosyl-(1->2)-alpha-D-Mannopyranosyl-(1->3)-[alpha-D-Mannopyranosyl-(1->6)-[alpha-D-Mannopyranosyl-(1->3)**]-**alpha-D-Mannopyranosyl-(1->6)]-beta-D-mannopyranosyl-(1->4)-N-acetyl-beta-D-glucopyranosyl-(1->4)**-**N-acetyl-beta-D-glucopyranosyl-1-OH
penta: alpha-D-Mannopyranosyl-(1->3)-[alpha-D-mannopyranosyl-(1->6)]-beta-D-mannopyranosyl-(1->4)-N-acetyl-beta-D-glucopyranosyl-(1->4)-N-acetyl-beta-D-glucopyranosyl-1-OH
PME: Particle Mesh Ewald
SNFG: Symbol Nomenclature for Glycans
TrpProAsn9glyco: H-D-Trp-L-Pro-D-Asn(-9glyco)-L-Pro-D-Trp-L-Pro-D-Asn-L-Pro-OH
TrpProAsnPenta: H-D-Trp-L-Pro-D-Asn(-Penta)-L-Pro-D-Trp-L-Pro-D-Asn(-Penta)-L-Pro-OH

### Appendix

#### Synthetic route

The molecules H-D-Trp-L-Pro-D-Asn(-Penta)-L-Pro-D-Trp-L-Pro-D-Asn(-Penta)-L-Pro-OH and H-D-Trp-L-Pro-D-Asn(-9glyco)-L-Pro-D-Trp-L-Pro-D-Asn-L-Pro-OH can be synthesized following these steps.

The peptide can be prepared manually by conventional solid phase chemistry on Fmoc-L-Pro-Wang resin according to published protocols^28^. Fmoc-Xaa-OH wil be used for couplings.

The Fmoc-Asparagine containing N-acetyilglucosamine can be prepared^29,30^ or purchased.

The resulting glycosylated Asn can be reacted with oligosaccharides by using glycosyltransferase making the sugar chain elongation possible^30,31^.

The test of the Arginine N-glycosylation and Arginine N-rhamnosylation can be obtained, using the protected α- and β-rhamnosylated arginine amino acids, following the procedures reported in 26 and 27 references.

The structures reported in fig. 1-3, 5-6, and 8 are obtained by RasWin v.2.7.5

This research was supported by the “Departments of Excellence-2018” Program (*Dipartimenti di Eccellenza*) of the Italian Ministry of Education, University and Research, DIBAF-Department of University of Tuscia, Project “Landscape 4.0 – food, wellbeing and environment”, through the provision of free computing time on NARTEN Cluster Facility, given by Prof. Nico Sanna.

## Supplementary material

The file pept_penta_T6AO.mpg shows the association of fig.8 in rotation.

The file pept9glyco_N262.mpg shows the association of fig.11 in rotation.

They have been obtained as follows. The program MoluCAD shows the rotation of the molecule imported in pdb format (https://www.kinematics.com/products/molucad.php). The image screen is recorded by means of the program CamStudio in avi format (https://camstudio.org/). The avi format is converted in mpg format by means of the online converter (https://www.online-convert.com/)

